# Stable integration of an optimized inducible promoter system enables spatiotemporal control of gene expression throughout avian development

**DOI:** 10.1101/2020.06.22.165704

**Authors:** Daniel Chu, An Nguyen, Spenser S. Smith, Zuzana Vavrušová, Richard A. Schneider

## Abstract

Precisely altering gene expression is critical for understanding molecular processes of embryogenesis. Although some tools exist for transgene misexpression in developing chick embryos, we have refined and advanced them by simplifying and optimizing constructs for spatiotemporal control. To maintain expression over the entire course of embryonic development we use an enhanced *piggyBac* transposon system that efficiently integrates sequences into the host genome. We also incorporate a DNA targeting sequence to direct plasmid translocation into the nucleus and a D4Z4 insulator sequence to prevent epigenetic silencing. We designed these constructs to minimize their size and maximize cellular uptake, and to simplify usage by placing all of the integrating sequences on a single plasmid. Following electroporation of stage HH8.5 embryos, our tetracycline-inducible promoter construct produces robust transgene expression in the presence of doxycycline at any point during embryonic development *in ovo* or in culture. Moreover, expression levels can be modulated by titrating doxycycline concentrations and spatial control can be achieved using beads or gels. Thus, we have generated a novel, sensitive, tunable, and stable inducible-promoter system for high-resolution gene manipulation *in vivo*.

## Introduction

For thousands of years, birds have been employed to understand how embryos grow, with some of the earliest documented observations made in ancient Greece by Hippocrates (circa 430 BCE) and especially by Aristotle (circa 350 BCE), who methodically recorded detailed empirical descriptions of developing chickens at various stages in the egg (Needham, 1934; Kember, 1971; Churchill, 1991; Aristotle and Thompson, 2001). Such early work not only helped establish developmental biology as a discipline, but also illustrates the fact that artificial incubation and widespread access to embryos have always been a highly advantageous asset of avian model systems (Lillie, 1952; Romanoff, 1960; Hamilton, 1965; Le Douarin and McLaren, 1984; Stern, 1994; Mason, 1999a; Stern, 2005; Jheon and Schneider, 2009).

The ease at which diverse embryos can be stage-matched (Hamburger and Hamilton, 1951; Hamilton, 1965; Ricklefs and Starck, 1998; Starck and Ricklefs, 1998; Schneider and Helms, 2003; Lwigale and Schneider, 2008; Ainsworth et al., 2010; Mitgutsch et al., 2011) combined with the commercial availability of fertilized eggs, the ability to “window” and reseal the egg shell, the comparatively large size of the embryo, and the capability of starting and arresting growth at any time during development have significantly advanced the use of birds for a broad range of experiments. Birds remain particularly applicable for questions that are best answered through microsurgical manipulations (e.g., tissue recombination, transplants, ablation, or extirpations), cell labeling and live imaging (e.g., fluorescent dyes and other agents, *ex ovo* culture, or immunochemical detection of engrafted cells), gain- and loss-of-function strategies (e.g., implantation of reagent-soaked beads, insertion of cell pellets, injection of biochemicals, infection with retroviruses, or electroporation of constructs) and other experimental approaches (Johnston, 1966; Noden, 1975; Serbedzija et al., 1989; Fekete and Cepko, 1993b; Stocker et al., 1993; Bronner-Fraser, 1996; Chen et al., 1999; Kulesa and Fraser, 2000; Larsen et al., 2001; Nakamura and Funahashi, 2001; Schneider et al., 2001; Garcia-Castro et al., 2002; Trainor et al., 2002; Cerny et al., 2004; Krull, 2004; Lwigale et al., 2004; Lwigale et al., 2005; Schneider, 2007; Bronner-Fraser and Garcia-Castro, 2008; Lwigale and Schneider, 2008; Sauka-Spengler and Barembaum, 2008; Fish and Schneider, 2014; Fish et al., 2014; Ealba et al., 2015; Woronowicz et al., 2018). Overall, such strategies have been indispensable to understanding numerous dynamic aspects of development including cell fate, tissue interactions, pattern formation, morphogenesis, and gene function and regulatory networks (Le Douarin and McLaren, 1984; Noden, 1984; Le Douarin et al., 1996; Clarke and Tickle, 1999; Schneider, 1999; Eames and Schneider, 2005; Noden and Schneider, 2006; Sauka-Spengler and Bronner-Fraser, 2008; Tokita and Schneider, 2009; Betancur et al., 2010; Le Douarin and Dieterlen-Lievre, 2013; Martik and Bronner, 2017; Abramyan and Richman, 2018; Schneider, 2018; Gammill et al., 2019; Nunez-Leon et al., 2019).

However, there are limitations to what can be done with avian embryos. First, the initial stages of development from the time of fertilization to the emergence of the blastoderm all happen before an egg is laid (Lillie, 1952; Romanoff, 1960; Kochav et al., 1980; Mason, 1999a). Thus, the earliest events are difficult to access and study. Moreover, by the time a fertilized egg is laid the embryo has already divided into around 60,000 cells (Mason, 1999b; Bellairs and Osmond, 2005). Second, avian eggs can be challenging to manipulate due to the size and abundance of the yolk. Third, unlike mouse or zebrafish model systems, birds have limited genetic tools (Hutt, 1949; Jull, 1952; Abbott, 1967) and targeted mutagenesis followed by forward genetics is difficult. While some transgenic chick and quail lines have been generated (McGrew et al., 2004; Chapman et al., 2005; Koo et al., 2006; van de Lavoir et al., 2006a; van de Lavoir et al., 2006b; Sato et al., 2010; Bower et al., 2011; Huss et al., 2015; June Byun et al., 2017; Tsujino et al., 2019), the technical challenges and expense of making transgenics, combined with the logistics of keeping sufficient transgenic flocks has limited the broad application of this approach (Sang, 2006). However, the ability to create genetic mutations through CRISPR/Cas9 technology has already made the prospects of genome engineering much easier in avians (Ahn et al., 2017; Gandhi et al., 2017; Morin et al., 2017; Williams et al., 2018).

Given the challenges of germ line transgenesis, proxies for studying gene function in avian model systems have predominantly involved a range of alternative strategies. For example, transgenes can be delivered efficiently using retroviral vectors (Fekete and Cepko, 1993a; Morgan and Fekete, 1996; Logan and Tabin, 1998; Chen et al., 1999; Kardon et al., 2003; Hughes, 2004) especially the replication-competent RCAS and RCASBP retroviruses. Some advantages of these vectors include their ability to spread widely throughout host tissues, which in turn allows for broad misexpression of a given transgene, and the ease at preparing large quantities of high-titer viral stocks (Logan and Tabin, 1998). But some limitations of retroviral vectors include the size of the gene insert that they can carry (up to approximately 2.4 kb), as well as their inability to infect most strains of chickens and other birds because of immunity arising from prior exposure to avian sarcoma-leukosis viruses (Hughes, 2004). A further draw-back of retroviral-based strategies is their general lack of precise control over the timing, spatial domains, and levels of gene misexpression. Oftentimes, to achieve sufficient amounts of viral spread, infection must be performed at very early stages, which means that the transgene has to be expressed continuously throughout development regardless if there is a specific stage desired for expression gene of interest.

Another approach for transiently misexpressing genes in a given location or for a certain period of time relies on electroporation of promoter-driven DNA constructs. Electroporation, which is very effective in avian embryos, involves placing electrodes to generate a pulsed electric field that transiently alters the plasma membrane and allows DNA constructs to be introduced into cells (Funahashi et al., 1999; Itasaki et al., 1999; Momose et al., 1999; Nakamura and Funahashi, 2001; Swartz et al., 2001; Chen et al., 2004; Krull, 2004; Simkin et al., 2014; Reberšek, 2017; McLennan and Kulesa, 2019). Several DNA constructs containing a robust chicken β-actin promoter, a CMV promoter, an internal ribosome entry site (IRES), and a bicistronic reporter with green fluorescent protein (GFP) have been widely adopted including pMES, pCIG, and pCAβ (Swartz et al., 2001; Megason and McMahon, 2002; McLarren et al., 2003; Sauka-Spengler and Barembaum, 2008; Jhingory et al., 2010; Hall et al., 2014; Yang et al., 2014; Gammill et al., 2019; Wu and Taneyhill, 2019). Electroporation can also efficiently enable gene repression using RNA interference (RNAi) and antisense morpholino oligonucleotides (Tucker, 2001; Kos et al., 2003; Chesnutt and Niswander, 2004; Krull, 2004; Nakamura et al., 2004; Rao et al., 2004; Das et al., 2006; Sauka-Spengler and Barembaum, 2008; Gammill et al., 2019). However, due to the extrachromosomal nature of these vectors such treatments are only transient since plasmids and short oligonucleotides degrade and dilute following the proliferation of transfected cells, and misexpression is almost entirely eliminated by 72 to 96 hours (Sauka-Spengler and Barembaum, 2008; Wang et al., 2011; Hall et al., 2014; Bourgeois et al., 2015). Moreover, the promoters in these widely used plasmids cannot be induced to control the timing or levels of gene expression. Thus, there has remained a need for highly versatile vectors that can achieve both long-term and conditional expression in avian embryos. To this end, one transgene expression system was created that uses *Tol2* transposon-mediated gene transfer (Koga et al., 1996) to enable stable integration of a given transgene into the avian genome (Kawakami, 2007), and that leverages a tetracycline (tet)-dependent inducible promoter (Sato et al., 2007; Watanabe et al., 2007; Takahashi et al., 2008). This system has been useful, for example, for studying the behavior and activity of neural crest mesenchyme (NCM) during later stages of embryogenesis (Yokota et al., 2011).

Building on the clear advantages of inducible promoter systems for exerting spatiotemporal control over gene expression and the ability of transposable elements to integrate into the avian genome and facilitate long-term expression throughout development (Wang et al., 2011; Macdonald et al., 2012; Serralbo et al., 2013; Bourgeois et al., 2015), we have designed a new gene delivery system that represents a substantial advance of this technology. Our goal was to streamline and minimize the number of components, to optimize the delivery and detection features, and to achieve efficient and more robust transgene expression. To do so, we generated a constitutively active mNeongreen (GFP) (Shaner et al., 2013) and doxycyline (dox)-inducible (Gossen et al., 1995; Loew et al., 2010; Heinz et al., 2011) mScarlet-I (RFP) (Bindels et al., 2017) construct. Then, to maintain expression of our electroporated constructs over the entire course of embryonic development, we combined our dox-inducible system with an enhanced *piggyBac* transposon system, which allows for stable semi-random integration into the host genome so that the construct is replicated along with the host genome (Lacoste et al., 2009; Lu et al., 2009; Yusa et al., 2011; Liu et al., 2013; Yusa, 2015). We further improved this construct by adding a D4Z4 genetic insulator sequence to block transcriptional repression (Bire et al., 2013) and a DNA targeting sequence (DTS) to direct transport of the plasmids into the nucleus (Dean et al., 1999; Bai et al., 2017). We find that this construct is sensitive to induction by dox both *in ovo* and in culture, integrates stably into the genome of chick and duck, and enables expression in embryonic tissues at any desired time or place. Here we demonstrate for example, that NCM can be electroporated at embryonic stage (HH) 8.5 and then gene expression can be induced at HH15, HH30, or later. Also, we show that transgene expression levels can be modulated by titrating the concentration of dox, and precise spatial control over transgene activation can be achieved by implanting beads or gels that release dox locally. Thus, our optimized and integrating inducible-promoter system can control the timing, spatial domains, and levels of gene misexpression throughout avian development, which will be useful for a broad range of experimental contexts.

## Results and Discussion

### Design of the small plasmid pNano

To maximize transfection and electroporation efficiency we aimed to generate plasmids as small as possible. Smaller plasmids have been shown to transfect and electroporate more efficiently than large plasmids (Yin et al., 2005). Moreover, large plasmids have been found to be toxic when introduced into cells independent of transgene expression from the plasmid (Lesueur et al., 2016). To minimize the size of our constructs we generated a new plasmid, pNano containing a minimal pMB1 plasmid origin of replication and β-lactamase resistance (BlaR) sequence with a small multicloning site containing EcoRI, EcoRV, and XhoI restriction enzyme sites. The plasmid is 1562 bp (Fig. 1A) and serves as the backbone for the other constructs generated. To our knowledge, pNano is the smallest plasmid with BlaR selection.

**Figure 1.**
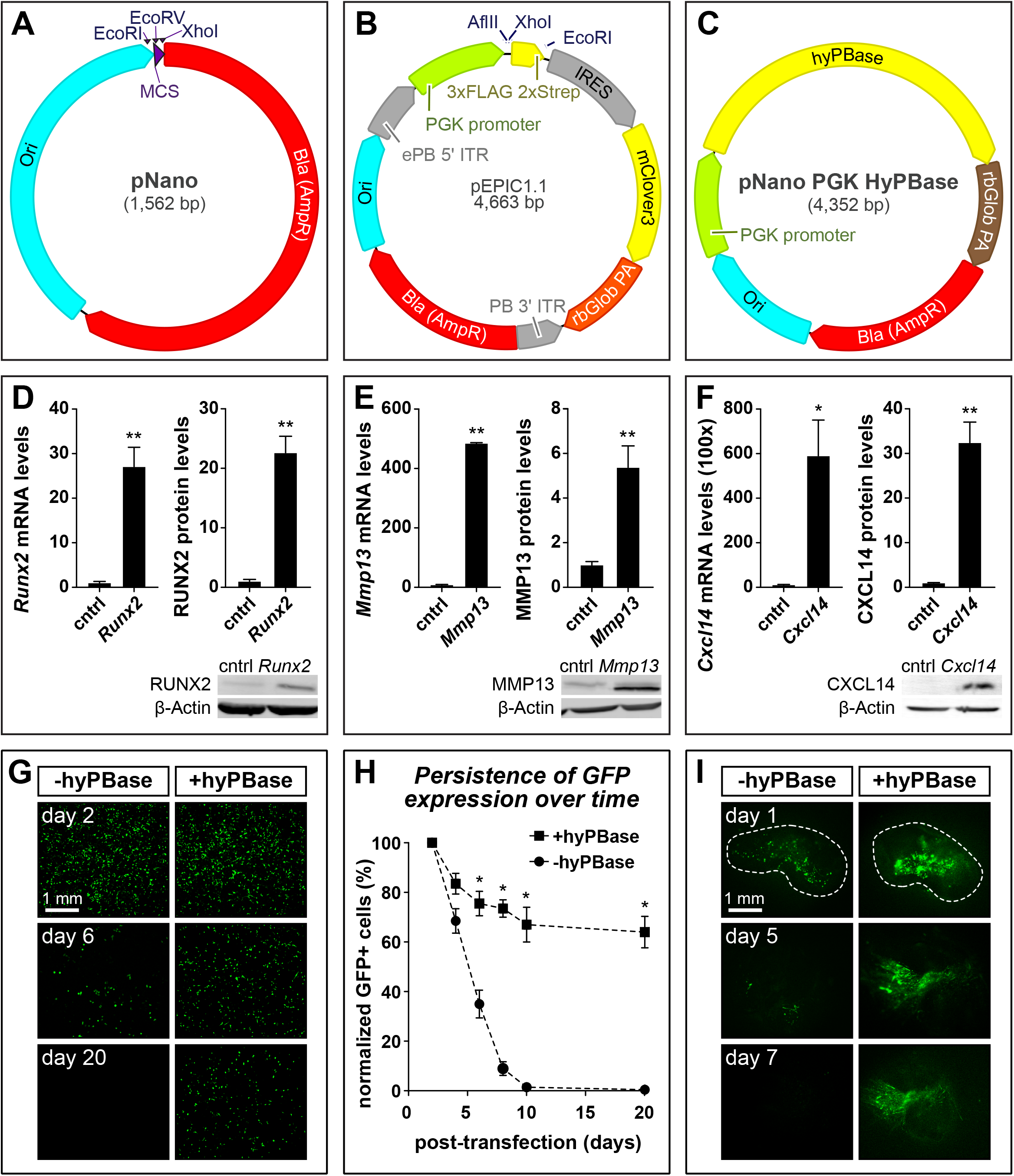
Plasmid maps and over-expression analyses. **A)** Map of the pNano minimal cloning vector showing restriction sites for cloning, multicloning sites (MCS) in purple, bacterial origin of replication (Ori) in cyan, and bacterial β-lactamase (Bla) resistance gene (AmpR) in red. **B)** Map of the pEPIC1.1 *piggyBac*-integrating constitutively-active expression vector showing *piggyBac* ITRs and IRES sequences in grey, PGK promoter sequences in green, terminator sequences in brown, and coding sequences in yellow. The pEPIC1.1 vector constitutively expresses mClover3, a green fluorescent protein (GFP). **C)** Map of the pNano-hyPBase expression vector used to integrate *piggyBac* sequences into host genome. **D)** Over-expression of *Runx2*, **E)** *Mmp13*, and **F)** *Cxcl14* with pEPIC1.1. DF-1 cells were transfected with control (cntrl) empty pEPIC1.1 or pEPIC1.1 plus *Runx2*, *Mmp13*, or *Cxcl14* coding sequences and harvested 3 days post-transfection. Relative mRNA levels were measured by qPCR and normalized using *18S*. Relative protein levels were measured by western blot (WB) and normalized using β-Actin. Representative WBs are shown below. N = 4 for *Runx2* and *Mmp13*, n = 2 for *Cxcl14*. **G)** Fluorescent images showing a time course of DF-1 cells transfected with pEPIC1.1. Cells were transfected either without pNano-hyPBase (left column) or with (right column). Cells were passaged every 2 days and imaged at 2, 6, and 20 days post-transfection. **H)** Quantification of GFP positive cells as a fraction of the total number of DF-1 cells transfected with pEPIC1.1 with or without pNano-hyPBase and normalized to 2 days post transfection. N = 2 for each group. **I)** Fluorescent images showing a time course of HH21 chick mandibular primordia electroporated with pEPIC1.1-*Cxcl14* either without pNano-hyPBase (left column) or with (right column) cultured, and imaged at day 1, 5, and 7. Error bars represent standard error of the mean (SEM). (*p < 0.05; **p < 0.005)

### Choosing a promoter

We chose the PGK1 promoter over other commonly used promoters for several reasons. The PGK1 promoter is relatively small at 500 bp and gives quite consistent expression across different cell types (Qin et al., 2010; Huss et al., 2015). The PGK1 promoter does not contain any viral sequences which are prone to epigenetic silencing and loss of expression over time (Brooks et al., 2004; Xia et al., 2007; Norrman et al., 2010).

### Choosing a transposon

Transient transfections and electroporations with standard plasmids only enable transgene overexpression for up to 5 days, which is much shorter than the time required to span *in ovo* development (e.g., 21 days for chick and 28 days for duck). To ensure stable and robust expression over the course of embryogenesis, we used a type II transposon (cut and paste) system to integrate sequences into the genome (Curcio and Derbyshire, 2003; Hickman et al., 2010; Yuan and Wessler, 2011). Several transposable systems currently exist including *Tol2* (Koga et al., 1996; Kawakami, 2007), *Sleeping Beauty* (Ivics et al., 1997), and *piggyBac* (Fraser et al., 1983; Fraser et al., 1996; Ding et al., 2005). We chose *piggyBac* due to its several advantages over other transposon systems. Most importantly *piggyBac* has been shown to have higher transposition activity than *Tol2* or *Sleeping Beauty* in human and chick (Wu et al., 2006; Lu et al., 2009; Huang et al., 2010) and there are improved versions of both the *piggyBac* transposon and transposase (Lacoste et al., 2009; Yusa et al., 2011). The efficiency of *piggyBac* integration is relatively size independent up to at least 10 kb (Ding et al., 2005) and *piggyBac* can deliver cargos in the hundreds of kb (Li et al., 2011; Rostovskaya et al., 2013), while *Sleeping Beauty* has reduced integration efficiency with cargo sizes above 5 kb (Geurts et al., 2003). *PiggyBac* semi-randomly integrates into genomes at sites of open chromatin while *Sleeping Beauty*’s integration pattern appears more random (Huang et al., 2010). Successful transposition events into silenced or heterochromatic regions may show no transgene expression due to epigenetic silencing. *PiggyBac* has lower rates of transgene silencing than *Sleeping Beauty* or *Tol2* (Meir and Wu, 2011). The *piggyBac* system is also relatively insensitive to the ratio of transposon to transposase while *Sleeping Beauty* and *Tol2* require titration to determine the optimal ratios (Meir et al., 2011). *PiggyBac* has consistent transposition activity across different cell lines (Wu et al., 2006) and has been utilized in many different organisms including yeast, mice, rats, humans, goat, pig, macaque, chick, rice, and several species of protists and insects (Yusa, 2015). This allows for the same construct to be used among different organisms compared to viral methods which have species-specificity.

### Generating the pEPIC1.1 construct for constitutive expression

To enable long term constitutive transgene expression, we generated pEPIC1.1 (<u>e</u>nhanced *<u>p</u>iggyBac* <u>I</u>RES m<u>C</u>lover3) (Fig. 1B). This construct drives transgene expression under the constitutive PGK promoter. To improve translational efficiency, we included a Kozak sequence directly upstream of the translational start site (Kozak, 1986). As a marker for expression, we used a minimal encephalomyocarditis virus internal ribosome entry site (IRES) (Bochkov and Palmenberg, 2006) to express a bicistronic transcript containing the overexpressed transgene and mClover3 (GFP) (Bajar et al., 2016). An optional C-terminal tandem affinity purification (TAP) tag consisting of 3xFLAG peptide sequences and 2xStrep-tag II sequences (Dalvai et al., 2015) can be added to enhance detection or pulldown. Sequences can be cloned either untagged by digesting pEPIC1.1 with AflII and EcoRI or tagged by digesting with AflII and XhoI. Sustained expression over long time periods is maintained by flanking the overexpression cassette with *piggyBac* inverted terminal repeat sequences (ITR). The ITRs in the presence of *piggyBac* transposase (PBase) semi-randomly integrates into the host genome at sequences containing a TTAA motif through a cut and paste mechanism. We used the enhanced *piggyBac* sequence which contains two point mutations in the left ‘5 ITR that increase transposition efficiency (Lacoste et al., 2009). To express PBase we also generated a complementary plasmid, pNano-hyPBase (Fig. 1C). This plasmid expresses a hyperactive version of PBase (hyPBase) (Yusa et al., 2011) under the PGK promoter.

As a proof-of-concept and to test expression of pEPIC1.1 we cloned coding sequences of a transcription factor (i.e., *Runx2*, 1419 bp), an extracellular matrix molecule (i.e., *Mmp13*, 1416 bp), and a cytokine (i.e., *Cxcl14*, 297 bp), into pEPIC1.1. We first confirmed that pEPIC1.1 constructs could overexpress our genes of interest by transfecting them into a chick fibroblast cell line (DF-1). We found that pEPIC1.1-*Runx2*, pEPIC1.1-*Mmp13*, and pEPIC1.1-*Cxcl14* all induce strong overexpression compared to empty pEPIC1.1 (Fig. 1D-F). The pEPIC1.1-*Runx2* construct increased *Runx2* mRNA levels 27 ±4.3 times by qPCR (p < 0.005) and RUNX2 protein levels 23 ±2.7 times by western blot (WB) compared to pEPIC1.1 (p < 0.005) (Fig. 1D). The pEPIC1.1-*Mmp13* construct increased *Mmp13* mRNA levels 480 ±2.4 times by qPCR (p < 0.005) and the MMP13 protein levels 5 ± 0.96 times by WB compared to pEPIC1.1 (p < 0.005) (Fig. 1E). The pEPIC1.1-*Cxcl14* construct increased *Cxcl14* mRNA levels 59,000 ±16,000 times by qPCR (p < 0.02) and the CXCL14 protein levels 32 ± 4.6 times by WB compared to pEPIC1.1 (p < 0.005) (Fig. 1F).

To confirm stable expression, we transfected DF-1 cells with pEPIC1.1 with or without pNano-hyPBase. Following transfection, cells were allowed to express GFP for 2 days to determine the baseline transfection efficiency. We then passaged the cells every 2 days for 20 days, to determine the stability of expression. We found that cells transfected without pNano-hyPBase rapidly lost GFP expression while those transfected with pNano-hyPBase initially had a small drop in GFP expression which then stabilized over time. At 6 days post transfection, cells with pNano-hyPBase retained higher levels of GFP expression compared to those without pNano-hyPBase (75% ±5 compared to 35% ±6, respectively, p < 0.05) (Fig. 1G-H). By day 20, 70% ±6 of cells transfected with pNano-hyPBase still expressed GFP, compared to < 1% of cells without pNano-hyPBase.

We next confirmed that the pEPIC1.1 construct is functional at the tissue level. Mandibular primordia (i.e., “mandibles”) were dissected from HH24 chick embryos, injected with a plasmid solution containing pEPIC1.1-*Cxcl14* with or without hyPBase, and then electroporated. Mandibles were then cultured over 7 days. After 5 days of culture, mandibles electroporated with pNano-hyPBase retained strong GFP expression while mandibles without pNano-hyPBase had greatly reduced expression compared to 1-day post-electroporation (Fig. 1I). After 7 days of culture mandibles electroporated without pNano-hyPBase had no detectable GFP expression.

### Generating the pPID2 piggyBac cloning vector

To enhance the versatility of our *piggyBac* vectors we generated a general *piggyBac* cloning vector pPID2 (*<u>p</u>iggyBac*, <u>i</u>nsulator, <u>D</u>TS) (Fig. 2A). pPID2 uses the pNano backbone to maintain a minimal vector footprint and contains the enhanced *piggyBac* mutations (Lacoste et al., 2009), a DNA nuclear targeting sequence (DTS), insulator sequence, and a multicloning site with over 20 restriction enzyme sites including HindIII, PstI, SalI, XhoI, EcorI, PstI, NcoI, NgoMV, NheI, SpeI, MscI, and BglII, for ease of cloning.

**Figure 2.**
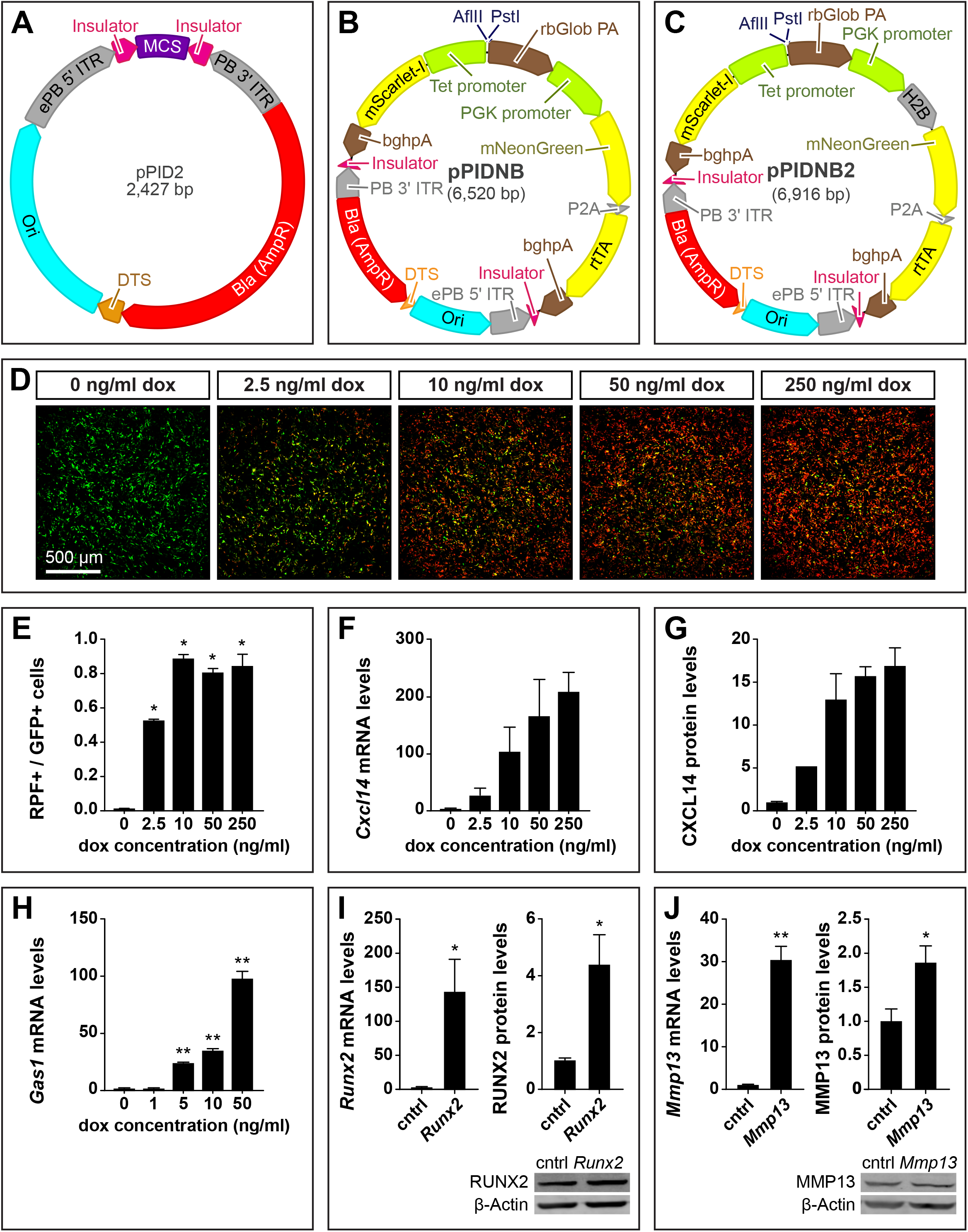
Maps of doxycycline (dox)-inducible plasmids and over-expression analyses. **A)** Map of the pPID2 *piggyBac* cloning vector showing insulators in magenta; a DNA nuclear targeting sequence (DTS) in orange; multicloning sites (MCS) in purple; bacterial origin of replication (Ori) in cyan; bacterial β-lactamase (Bla) resistance gene (AmpR) in red; and *piggyBac* ITRs, IRES, and P2A sequences in grey. **B)** Map of the pPIDNB *piggyBac* dox-inducible vector showing restriction sites for cloning, coding sequences in yellow, terminator sequences in brown, and promoter sequences in green. pPIDNB constitutively expresses mNeonGreen (GFP) and coding sequences can be cloned into the plasmid under a bidirectional tetracycline (tet) inducible promoter using the AflII and PstI restriction sites. mScarlet-I, a red fluorescent protein (RFP), is expressed on the alternate side of the bidirectional tet promoter. **C)** Map of the pPIDNB2 vector, which is identical to pPIDNB except that GFP is localized to the nucleus using histone H2B. **D)** DF-1 cells transfected with pPIDNB constitutively express GFP and differentially express RFP in response to varying concentrations of dox after 24 hours. **E)** RFP-positive (i.e., dox-induced) cells relative to total number of GFP-positive (i.e., transfected) cells. **F)** dox induction was measured in DF-1 cells on the mRNA and **G)** protein levels for *Cxcl14* (n = 2 for each group except n = 1 for the 2.5 ng/ml treatment; and mRNA level for **H)** *Gas1* (n = 4 for each group). Levels are relative to 0 ng/ml of dox and normalized to *18S* for mRNA and β-Actin for protein. **I)** Over-expression of *Runx2* and **J)** *Mmp13* with pPIDNB. DF-1 cells were transfected with control (cntrl) empty pPIDNB or pPIDNB plus *Runx2* or *Mmp13* coding sequence and treated with 50 ng/ml of dox for 24 hours. mRNA levels were normalized using *18S* and protein using β-Actin. Representative WBs are shown below. N = 4 for each group. Error bars represent SEM. (*p < 0.05; **p < 0.005)

When cells are transfected or electroporated with plasmids, transport from the cytoplasm to the nucleus is required for both expression and transposition into the genome. Plasmid entry into the nucleus generally occurs either during mitosis when the nuclear envelope breaks down, allowing for passive diffusion of plasmids into the nuclear space, or when the intracellular plasmid concentration is very high (10^4^ – 10^6^ molecules of plasmid DNA per cell) (Utvik et al., 1999; Young et al., 2003; Bai et al., 2017). To overcome potential nuclear import barriers, we added a DTS (Dean et al., 1999; Dean et al., 2005). DTSs function by binding to transcription factors which are then actively transported into the nucleus. We chose to use the simian virus 40 (SV40) 72 bp promoter DTS (Dean et al., 1999) because it can function in a wide variety of cell types (Dean, 1997; Young et al., 2003), is small, uses endogenously expressed transcription factors (Miller et al., 2009), and does not require expression of viral proteins (Dean et al., 2005). Alternatively, if nuclear entry is low even with a DTS, addition of trans-cyclohexane-1,2-diol reversibly increases the permeability of the nuclear pore complex allowing plasmids to diffuse into the nucleus (Vandenbroucke et al., 2007; De la Rossa and Jabaudon, 2015; Cervia et al., 2018).

Epigenetic and heterochromatic silencing of foreign DNA inserted into the host genome represent an obstacle for efficient transgene expression both at the time of insertion and over long-term expression (Garrison et al., 2007). Genomic insertions containing viral sequences are known to be actively silenced (Pannell and Ellis, 2001; Ellis, 2005; Wen et al., 2014; Hudecek et al., 2017). An insertion in a heterochromatic region or region that subsequently becomes heterochromatic may result in transgene inactivation (Janssen et al., 2018). To prevent this epigenetic silencing, we added a genetic insulator that blocks the spread of repressive epigenetic marks and heterochromatin (Ali et al., 2016). Moreover, insulator sequences help to protect endogenous sequences from epigenetic activation or silencing caused by the transposition (Hollister and Gaut, 2009). We used the D4Z4 insulator, which is only 65 bp and has been shown to efficiently protect *piggyBac* transgene expression (Ottaviani et al., 2009; Bire et al., 2013). pPID2 contains two D4Z4 insulator sequences contained within the *piggyBac* ITRs flanking the multicloning site (Fig. 2A).

### Generating the pPIDNB doxycycline-inducible system

We also added a dox-inducible component to our overexpression constructs, which provides several advantages, including increased temporal control of expression. Without such precise temporal control, the premature and continuous expression of a gene of interest may disrupt development in ways that cause phenotypes unrelated to the processes under study. A dox-based strategy has several advantages over other inducible systems in that dox is extremely cheap and effective at low concentrations. Additionally, dox is able to diffuse efficiently through tissues allowing for induction past the surface level (Agwuh and MacGowan, 2006; Sato et al., 2007) and the use of dox-soaked beads or gels can allow for spatial control of expression.

We generated the pPIDNB (*<u>p</u>iggyBac*, insulator, <u>D</u>TS, m<u>N</u>eongreen, <u>b</u>i-directional) construct as a minimal dox-inducible plasmid (Fig. 2B). This plasmid is based upon the pPID2 backbone and includes the DTS, insulator, and *piggyBac* sequences. In addition, pPIDNB constitutively expresses the reverse tetracycline (tet) transactivator (rtTA) and mNeongreen (GFP) under the PGK promoter (Shaner et al., 2013). The rtTA and mNeongreen coding sequences are bicistronic and are separated by a porcine teschovirus-1 2A (P2A) site which causes them to be expressed as two different peptide sequences (Szymczak et al., 2004; Kim et al., 2011). When bound to dox, the rtTA undergoes a conformational shift allowing binding and activation of the bidirectional tet promoter (Gossen et al., 1995; Das et al., 2016). We chose to use the rtTA-V16 variant of rtTA, which is both sensitive to dox and can induce strong expression (Das et al., 2016). On one side of the bidirectional promoter is mScarlet-I (RFP) serving as a marker for dox induction (Bindels et al., 2017). On the other side of the bidirectional promoter is the cloning site containing AflII and PsiI sites for dox-inducible expression of the gene of interest. Combining both the rtTA and tet promoter into a single construct enables stable inducible-expression with one integrating plasmid and one transposase-expressing plasmid. Moreover, for experiments that would benefit from the ability to detect nuclear localization, we also generated pPIDNB2, which has histone H2B fused to GFP to label nuclei, in contrast to the pPIDNB plasmid where GFP localization is diffuse throughout the cell (Fig. 2C).

To evaluate the sensitivity of the pPIDNB plasmid to induction by dox, we transfected DF-1 cells and performed a dose-response analysis with dox for 24 hours. In the absence of dox, there was a very low basal level of RFP expression, with only 0.15%±0.2% of the GFP positive cells also expressing detectable levels of RFP expression (Fig. 2D-E). After treating cells with 2.5 ng/ml dox, 52% ±1.1% of the GFP positive cells also expressed RFP. We found that the percent of RFP expressing cells as a fraction of the GFP positive cells maxed out at a dose of 10 ng/ml dox at 88% ±2.7% with cells treated at 50 ng/ml and 250 ng/ml dox expressing RFP at 80% ±2.7% and 84% ±7.3%, respectively (Fig. 2E). While the fraction of cells expressing RFP did not increase at dox concentrations greater that 10 ng/ml, the intensity of RFP did increase with higher concentrations of dox (Fig. 2D).

We next tested the ability of pPIDNB to drive exogenous gene expression by cloning in the coding sequences for *Cxcl14*, *Gas1* (a plasma membrane receptor, 945 bp), *Runx2*, and *Mmp13*. We first transfected DF-1 cells with pPIDNB-*Cxcl14*, treated with various doses of dox, and found that *Cxcl14* expression correlated with the concentration of dox (Fig 2F). These results along with the RFP data above indicate that dox dose-response is tunable at the cellular level and not simply a binary response to increased dox concentrations causing more cells to express RFP. We found DF-1 cells treated with 2.5, 10, 50, and 250 ng/ml dox for 24 hours increased *Cxcl14* mRNA expression by 26 ±14, 96 ±44, 150±70, and 160 ±55 times respectively, compared to cells not treated with dox (Fig. 2F). WB analysis also showed a dose response with 2.5, 10, 50, and 250 ng/ml dox with CXCL14 protein levels increasing by 5.2, 13 ±3.1, 16 ±1.2 and 17 ±2.2 times respectively, compared to cells not treated with dox (Fig. 2G).

To determine if pPIDNB can stably integrate into the genome and express a transgene, we transfected DF-1 cells with pPIDNB-*Gas1* and pNano-hyPBase. DF-1 cells were passaged over 4 weeks and then fluorescence-activated cell sorted (FACS) for GFP to confirm pPIDNB-*Gas1* could be stably integrated into the host genome and remain dox-inducible. We treated cells with dox and found that they were induced in a dose-response manner. After treating cells with 1, 5, 10, and 50 ng/ml dox for 24 hours, *Gas1* mRNA expression increased by 1.1 ±0.16 (p > 0.05), 23 ±0.99 (p < 0.005), 34 ±2.7 (p < 0.005), and 97 ±7.6 times (p < 0.005), respectively, compared to cells not treated with dox (Fig. 2H). To confirm that pPIDNB can overexpress different types of genes we also transfected DF-1 cells with either with empty pPIDNB, pPIDNB-*Runx2*, or pPIDNB-*Mmp13*. Transfected cells were treated with 50 ng/ml of dox for 24 hours. The pPIDNB-*Runx2* and pPIDNB*-Mmp13* transfected cells expressed 140 ±47 (p < 0.05) times more *Runx2* mRNA and 30 ±3.2 (p < 0.005) times more *Mmp13* mRNA than cells transfected with empty pPIDNB, respectively (Fig. 2I-J). WB analyses also showed overexpression with pPIDNB-*Runx2* and pPIDNB-M*mp13* expressing 4.4 ±1.1 (p < 0.05) and 1.9 ±0.25 (p < 0.05) times more RUNX2 and MMP13 protein than pPIDNB alone, respectively.

### Spatiotemporal control of expression in cell culture

To confirm that we could exert spatiotemporal control over transgene expression using pPIDNB, DF-1 cells were transfected with pNano-hyPBase and either pPIDNB-*Gas1* or pPIDNB2-*Gas1*. Cells were passaged for 4 weeks and then sorted for GFP to generate stable lines with either pPIDNB-*Gas1* or pPIDNB2-*Gas1* integrated into their genomes. Cells with integrated pPIDNB-*Gas1* or pPIDNB2-*Gas1* were visualized by GFP. pPIDNB-*Gas1* cells showed GFP localized throughout the entire cell while pPIDNB2-*Gas1* showed nuclear localization of GFP (Fig. 3A-B). Cells were then treated with 50 ng/ml dox and imaged at 0, 6, and 12 hours post dox treatment. After 6 hours of dox treatment, cells began to express detectable levels of RFP and by 12 hours the RFP signal was robust.

**Figure 3.**
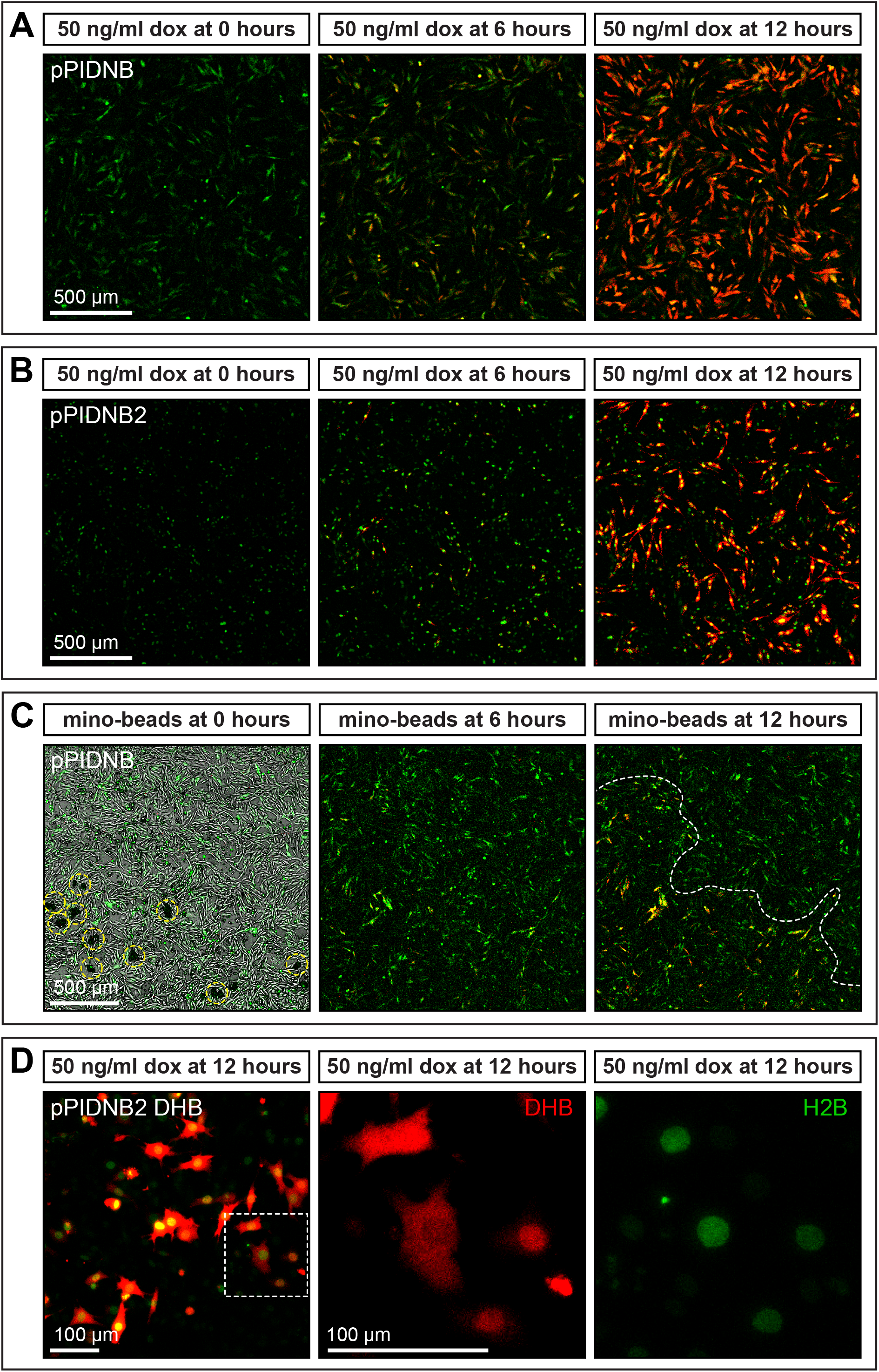
In vitro induction of gene expression in cells. **A)** DF-1 cells stably transfected with pPIDNB constitutively express mNeonGreen (GFP) and begin to express mScarlet-I (RFP) over time in response to treatment with 50 ng/ml doxycycline (dox). Cells were imaged at 0, 6, and 12 hours post treatment. **B)** DF-1 cells stably transfected with pPIDNB2 constitutively express GFP in the nucleus and begin to express RFP over time in response to treatment with 50 ng/ml dox. Cells were imaged at 0, 6, and 12 hours post treatment. **C)** DF-1 cells transfected with pPIDNB constitutively express GFP and begin to express RFP over time in response to treatment with minocycline microspheres. Cells were imaged at 0, 6, and 12 hours post treatment. Microspheres are circled in yellow and a boundary between cells that are induced versus those that are not is indicated by a white dashed line. **D)** DF-1 cells transfected with pPIDNB2 DHB (DNA Helicase B) constitutively express GFP in the nucleus and the DHB cell cycle sensor is tagged with RFP and induced in response to 50 ng/ml dox as seen at 12 hours post treatment. White-dashed inset box indicates cells shown at higher magnification where RFP marks DHB localization and GFP marks nuclei. DHB localization appears enriched in the nucleus, cytoplasm, or diffused throughout the cell.

To determine if we could control the spatial localization of transgene expression, we applied minocycline microspheres to DF-1 cells transfected with pPIDNB-*Gas1*. These microspheres slowly release minocycline, a tetracycline (dox) analog, and induce the tet expression system (Chtarto et al., 2003; Zhou et al., 2006). We applied minocycline microspheres directly to a localized area in the well and cells were imaged at 0, 6, and 12 hours after treatment. After 6 hours, we observed low levels of RFP expression, and after 12 hours RFP expression levels were high in areas adjacent to the microspheres but not in areas further away (Fig. 3C).

For experiments that could benefit from the ability to monitor dynamic changes in the cell cycle, we added a DNA helicase B (DHB) cell cycle sensor sequence (Spencer et al., 2013; Kohrman et al., 2019) to the dox-inducible RFP of pPIDNB2. The DHB cell cycle sensor translocates to the nucleus at G0/G1. During S phase, DHB localizes to both the nucleus and the cytoplasm and during M-phase DHB primarily localizes to the cytoplasm. The nuclear localization of GFP in pPIDNB2 allows for the determination of how much of the DHB signal is nuclear versus cytoplasmic. We transfected DF-1 cells with pPIDNB2 DHB and treated them with 50 ng/ml of dox and imaged them after 12 hours. We found that we could identify cells in different phases of the cell cycle with nuclear-localized DHB (G0/G1), nuclear- and cytoplasm-localized DHB (S phase), and cytoplasm localized DHB (M phase) (Fig. 3D).

### Temporal and spatial control of gene expression during development

To exert spatiotemporal control over gene expression in embryonic tissues, we unilaterally electroporated the premigratory cephalic NCM of HH8.5 chick embryos with pPIDNB and pNano-hyPBase. At HH10, we assayed for the extent of electroporation by visualizing GFP-positive cells *in ovo* in migrating NCM destined for the mandibular primordia (Fig. 4A). These embryos were then incubated until HH30, at which point the mandibular primordia were dissected out, cultured with 50 ng/ml of dox, and imaged at 0, 12, and 24 hours post-treatment. As evidence of the stable genomic integration and induction of the plasmids in embryos, we observe the electroporated side of the mandible expressing GFP, with the contralateral side showing little to no GFP expression. After 12 hours, treatment with dox results in strong RFP signal that is co-localized with GFP and this RFP expression intensifies further by 24 hours (Fig. 4A, Movie 1).

**Figure 4.**
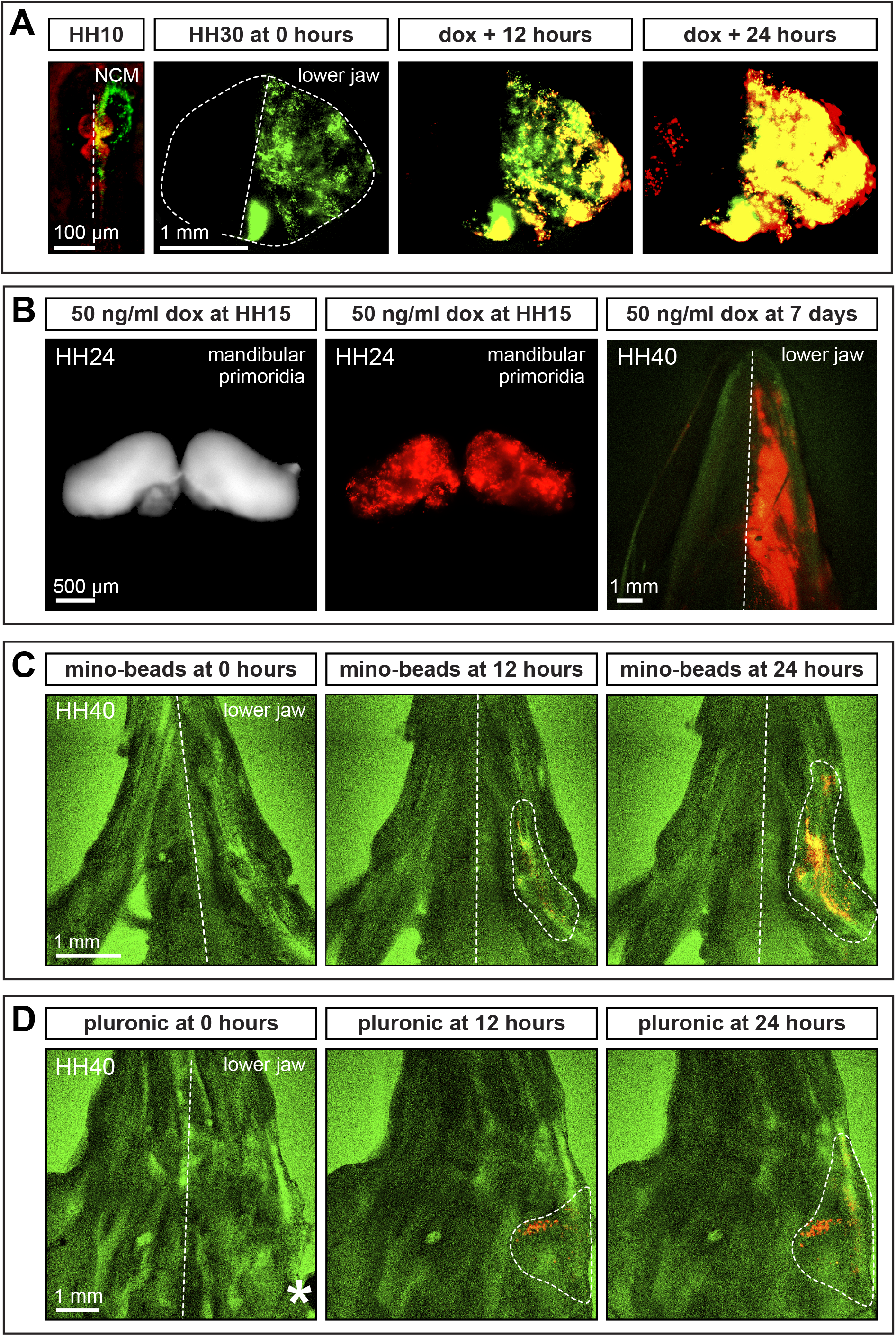
*In ovo* and *ex vivo* induction of gene expression in the lower jaw. **A)** Cephalic neural crest mesenchyme (NCM) electroporated unilaterally with pPIDNB and pNano-hyPBase in a chick embryo at HH8.5 constitutively expresses mNeonGreen (GFP) as shown at HH10 (counterstained red with neutral red). At HH30, the lower jaw shows unilateral GFP expression in NCM. After 12 and 24 hours in culture, NCM express mScarlet (RFP) in response to treatment with 50 ng/ml doxycycline (dox). **B)** NCM bilaterally electroporated with pPIDNB and pNano-hyPBase in a duck embryo at HH8.5 shows RFP expression on both sides of the lower jaw at HH24 (after bilateral electroporation) when treated *in ovo* with 50 ng/ml dox at HH15. NCM bilaterally electroporated with pPIDNB and pNano-hyPBase in a chick embryo at HH8.5 shows RFP expression on one side of the lower jaw at HH40 (after unilateral electroporation) when treated *in ovo* with 50 ng/ml dox 7 days after electroporation and imaged 9 days after treatment. **C)** NCM electroporated bilaterally with pPIDNB-*Gas1* and pNano-hyPBase in a duck embryo at HH8.5 shows RFP expression in the lower jaw (white dashed area) 12 and 24 hours after being injected in culture with minocycline microspheres at HH40. **D)** NCM electroporated unilaterally with pPIDNB-*Gas1* and pNano-hyPBase in a duck embryo at HH8.5 shows RFP expression in the lower jaw (white dashed area) 12 and 24 hours after being treated at HH40 with 35% Pluronic F-127 gel containing 50 ng/ml dox (white asterisk).

Additionally, some duck embryos were bilaterally electroporated at HH8.5 with pPIDNB*-Gas1* and pNano-hyPBase and were treated with 50 ng/ml dox *in ovo* at HH15. By HH24, we observed RFP expression throughout the mandibular primordia (Fig. 4B). To confirm that *in ovo* dox treatment would work efficiently even during later stages of development, some chick embryos were unilaterally electroporated at HH8.5 with pPIDNB and pNano-hyPBase, incubated for 7 days, and then were treated *in ovo* with a single dose of dox (50 ng/ml). These embryos were then allowed to develop for 9 more days (to around HH40), at which point we observed robust unilateral RFP expression in the lower jaw (Fig. 4B).

To exert more precise spatial control over gene expression, some embryos were bilaterally electroporated at HH8.5 with pPIDNB*-Gas1* and pNano-hyPBase and incubated until HH40. Their lower jaws were then harvested and either injected along the right side with minocycline microspheres or with dox gel (Pluronic F-127). Pluronic F-127 is a liquid at low temperatures (4 °C) but solidifies at higher temperatures (37 °C) and has been used for delivering drugs to different tissues (Harris et al., 2004; Giovagnoli et al., 2010). After 12 hours of treatment with either minocycline microspheres or with dox gel, we observe RFP expression localized on the right side of the jaw, which becomes more elevated by 24 hours (Fig. 4C-D).

## Conclusion

In this study we generated an “all-in-one” *piggyBac* dox-inducible system. The pPIDNB plasmid is designed to be as small as possible to optimize cellular uptake while incorporating critical features to maximize its functionality. The DTS and insulator sequences serve to promote expression by directing nuclear entry of the plasmid and block heterochromatic silencing expression. We used mutated *piggyBac* and hyPBase sequences to increase genome integration efficiency. We have also incorporated a constitutively expressed GFP to mark cells that have taken in plasmid DNA and RFP to mark dox-induced cells. Our system facilitates precise temporal control of gene induction and is easily adapted for *in vitro* or *in ovo*. Spatial control of gene expression can be achieved by electroporating regions of interest and/or by applying beads or gels to localize the distribution of dox.

The pPIDNB system is able to induce expression quickly and its reliance on a low dose of dox is important because dox has biological effects beyond antimicrobial activity including affecting matrix metalloproteinase activity, inflammation, the NF-κB pathway, and the nervous system (Bahrami et al., 2012; Alexander-Savino et al., 2016). High concentrations of dox (e.g., 1000 ng/ml) are cytotoxic in culture and have strong proliferative and metabolic effects, and some cell types are affected at even lower concentrations (e.g., 100-200 ng/ml) (Ermak et al., 2003; Ahler et al., 2013; Alexander-Savino et al., 2016). By using a low dose of dox (50 ng/ml) we have likely minimized any off-target effects of dox treatment.

While in the current study, we designed the pPIDNB construct for transgene overexpression, we envision that future applications will include different types of experiments such as gene knockdown using CRISPRi (Qi et al., 2013; Mandegar et al., 2016). For example, catalytically inactive *Cas9* could be placed with transcriptional repressors under an inducible tet promoter (Qi et al., 2013; Yeo et al., 2018). Constitutively active U6 promoters would drive expression of single guide RNAs (Cong et al., 2013; Gandhi et al., 2017; Williams et al., 2018). Using similar protocols for overexpression or knockdown would reduce the number of variables between experiments and help limit the confounding effects from different constructs. Overall, a great strength of avian model systems has been the combination of experimental embryology and modern genetic techniques. Our new, sensitive, stable, and robust inducible-promoter system builds on this strength and joins an arsenal of tools for manipulating gene expression that will likely be useful to the broader developmental biology community.

## Materials and Methods

### Plasmids

To generate pNano, the Ori and BlaR from pJet1.2 (ThermoFisher, K1231) were amplified using Q5 Hot Start High-Fidelity DNA Polymerase (NEB, M0493L). Fragments were cloned together using NEBuilder HiFi DNA Assembly Master Mix (NEB, E2621L). EcoRI, XhoI, and EcorV restriction enzyme sites were incorporated as tails added to the primers. To generate pEPIC1.1, the enhanced *piggyBac* ITRs, PGK promoter, 3x FLAG 2x Strep tag, IRES, mClover3, rabbit Beta globin terminator sequence, pNano were amplified by PCR using Q5 Hot Start High-Fidelity DNA Polymerase and cloned together using NEBuilder HiFi DNA Assembly Master Mix. The enhanced *piggyBac* ITRs were ordered as gBlocks (IDT). The 3x FLAG 2x Strep tag sequence was amplified from AAVS1 Puro Tet3G 3xFLAG Twin Strep (Addgene, # 92099) (Dalvai et al., 2015). mClover3 sequence was amplified from pKanCMV-mClover3-mRuby3 (Addgene, #74252) (Bajar et al., 2016). To generate pNano-hyPBase, the PGK promoter, hyPBase, and rabbit β-Globlin poly A sequences were amplified by PCR using Q5 Hot Start High-Fidelity DNA Polymerase and cloned together using NEBuilder HiFi DNA Assembly Master Mix. To generate pPID2, the SV40 72 bp DTS and two 65 bp insulator sequences flanking MCS were ordered as gBlocks (IDT). The enhanced *piggyBac* ITRs, Ori, and BlaR were amplified using Q5 Hot Start High-Fidelity DNA Polymerase and cloned together with the DTS and insulator gBlocks using NEBuilder HiFi DNA Assembly Master Mix. To generate pPIDNB, the bovine growth hormone poly A, mScarlet-I, bidirectional tet promoter, rabbit β-Globlin poly A, PGK promoter, mNeongreen P2A, and rtTA sequences were amplified by PCR using Q5 Hot Start High-Fidelity DNA Polymerase and then cloned together using NEBuilder HiFi DNA Assembly Master Mix. The bidirectional tet promoter and rtTA sequences were amplified from AAVS1 Puro Tet3G 3xFLAG Twin Strep (Addgene, # 92099). The mScarlet-I sequence was amplified from pmScarlet-i_C1 (Addgene, #85044) (Bindels et al., 2017). To generate pPIDNB2, H2B was amplified using Q5 Hot Start High-Fidelity DNA Polymerase and then cloned into pPIDNB with QuikChange (Liu and Naismith, 2008) using KOD Xtreme Hot Start DNA Polymerase (Sigma, 71975-3). To generate pPIDNB2-DHB, DHB was ordered as a gBlock and cloned into pPIDNB2 digested with XhoI (NEB, R0146S) and NotI (NEB, R3189S).

### RNA extractions

For *Runx2*, *Mmp13*, and *Cxcl14,* RNA was extracted from DF-1 cells and HH27 whole chick heads using the RNeasy Plus Kit (Qiagen, 74136) following the manufacturer’s directions. Whole heads and DF-1 cells were resuspended in 600μl of RTL plus buffer supplemented with 1% β-mercaptoethanol. Homogenization was carried out in a Bead Mill 24 (ThermoFisher, 15-340-163) at 5 m/s for 30 s. Following purification of total RNA, residual genomic DNA was removed using TURBO DNA-free Kit (Invitrogen, AM1907). For RNA extractions involving *Gas1*, The PicoPure RNA Isolation Kit (Applied Biosystems, KIT0204) was used following the manufacturer’s directions and homogenization was carried out in a Bead Mill 24 at 4 m/s for 15 s.

### Cloning coding sequences

Full length cDNA synthesis from RNA was carried out using Maxima H-reverse transcriptase (ThermoFisher, K1651) following the manufacturer’s directions with 2 μg of total RNA and 100 pmol of d(T)20 VN primer. The cDNA synthesis reaction was carried out at 50 °C for 30 min, 55 °C for 10 min, 60 °C for 10 min, 65 °C for 10 min, and 85 °C for 5 min. Full length *Runx2*, *Mmp13*, *Cxcl14*, and *Gas1* were amplified by PCR using Q5 Hot Start High-Fidelity DNA Polymerase (NEB, M0493L) and cloned using CloneJET PCR Cloning Kit (ThermoFisher, K1231). Following confirmation of cloning of full length coding sequences by Sanger sequencing, *Runx2*, *Mmp13*, *Cxcl14*, and *Gas1* were cloned into pEPIC1.1 digested with AflII (NEB, R0520S) and EcoRI (NEB, R3101S) or pPIDNB digested with AflII (NEB, R0520S) and PstI (NEB, R3140S) using NEBuilder HiFi DNA Assembly Master Mix. All constructs were verified by Sanger sequencing and midipreped for electroporation and transfection using PureLink Fast Low-Endotoxin Midi Kit (Invitrogen, A36227).

### Avian embryos and cell culture

Fertilized eggs of chicken (*Gallus gallus*) and duck (*Anas platyrhynchos*) were purchased from AA Lab Eggs (Westminster, CA) and incubated at 37.5 °C in a humidified chamber (GQF Hova-Bator, Savannah, GA, 1588) until they reached embryonic stages appropriate for manipulation and/or analyses. For all experiments, we adhered to accepted practices for the humane treatment of avian embryos as described in S3.4.4 of the AVMA Guidelines for the Euthanasia of Animals: 2013 Edition (Leary et al., 2013). Embryos were matched at equivalent stages using the Hamburger and Hamilton (HH) staging system, a well-established standard which utilizes an approach based on external morphological characters and that is independent of body size and incubation time (Hamburger and Hamilton, 1951; Hamilton, 1965; Ricklefs and Starck, 1998; Starck and Ricklefs, 1998). For late embryonic stages, we relied primarily on growth of the limbs, facial primordia, feather buds, and eyes (Eames and Schneider, 2005; Eames and Schneider, 2008; Merrill et al., 2008).

Embryonic chick fibroblasts (DF-1) were purchased (ATCC, CRL-12203) and cultured in Dulbecco’s Modified Eagle’s Medium (DMEM, Corning, 10-013-CV) supplemented with 10% FBS (VWR, 97068-085, Lot# 283K18) and 1X penicillin-streptomycin (ThermoFisher, 15140122) at 37 °C with 5% CO_2_. Cells were passaged twice a week. Cells were transfected with lipofectamine 3000 (ThermoFisher, L3000008) according to the manufacturer’s instructions. Transfections for integrating *piggyBac* vectors were carried out in 6-well plates with 5 μg *piggyBac* plasmid, 5 μg of pNano-hyPBase, and 20 ul of P3000.

### Electroporations

Electroporations were performed by injecting a solution of pEPIC1.1-Cxcl14 and pNano-hyPBase at 3 μg/μl and 1 μg/μl, respectively, with a small amount of Fast Green dye. DNA was injected with a Pneumatic PicoPump (PV830, World Precision Instruments) into dissected HH21 mandibular primordia using thin wall borosilicate glass micropipettes (O.D. 1.0 mm, I.D. 0.75 mm, Sutter Instrument, B100-75-10) pulled on a micropipette puller (P-97 Flaming/Brown, Sutter Instrument). Mandibles were placed between two gold plate electrodes 0.5 cm apart submerged in Hanks’ balanced salt solution (Sigma-Aldrich). Electroporations were carried out by delivering five square pulses at 25 V for 50 ms spaced 500 ms apart (CUY21EDITII Next Generation Electroporator, BEX CO, Ltd). Mandibles were then cultured in BgJB medium (ThermoFisher, 12591038) supplemented with 10% FBS (VWR, 97068-085, Lot# 283K18) and 1 X penicillin-streptomycin (ThermoFisher, 15140122).

*In ovo* electroporations were performed using a solution of pPIDNB and pNano-hyPBase at 3 μg/μl and 1 μg/μl, respectively. With the addition of Fast Green tracer dye, DNA solution was injected into HH8.5 chick neural tubes with a Pneumatic PicoPump using thin wall borosilicate glass micropipettes pulled on a micropipette puller. Platinum electrodes were positioned on each side of the area pellucida, centered on the midbrain-hindbrain boundary. For unilateral electroporations, we delivered three square pulses at 50 V for 1 ms spaced 50 ms apart followed by five square pluses at 10 V for 50 ms spaced 50 ms apart. For bilateral electroporations, we delivered three square pulses at 50 V for 1 ms spaced 50 ms apart, three square pulses at 50 V for 1 ms spaced 50 ms apart in the reverse polarity, three five square pluses at 10 V for 50 ms spaced 50 ms apart followed by, five square pluses at 10 V for 50 ms spaced 50 ms apart in the reverse polarity.

### qPCR

DNased RNA was reverse-transcribed using iSCRIPT (Bio-Rad, 1708841). Gene expression was quantified by qPCR with iQ SYBR Green Supermix (Bio-Rad, 1708882) and normalized to 18S rRNA following previously published protocols (Dole et al., 2015; Smith et al., 2016). Primer sets were designed and optimized as described previously (Ealba and Schneider, 2013) and are listed in supplemental Table S1. Each sample was assayed in technical duplicate.

### Western blot

DF-1 cells were lysed with 1X RIPA lysis buffer (EMD Millipore, 20-188) containing Halt protease inhibitors (ThermoFisher, 78430). A BCA assay (ThermoFisher, 23225) using a SpectraMax M5 plate reader was performed to quantify protein, and 40 μg protein was electrophoresed on a 10% SDS polyacrylamide gel following a published protocol (Smith et al., 2016). Proteins were transferred to an Imobilon-PPVDF membrane (Millipore, Billerica, MA, IPVH00010). Membranes were probed with rabbit anti-chick RUNX2 primary antibody (1:1000, AbCam Burlingame, CA, Cat #ab23981), custom made rabbit anti-chick MMP13 antibody (1μg/ml, Genscript), rabbit anti-CXCL14 (0.2 μg/ml, Peprotech, 500-P237), mouse anti-chick β-actin antibody (1:4000, Novus Biologicals, NB600-501), goat anti-rabbit IRDye 800CW (1:15000, LI-COR #925-32211), and donkey anti-mouse IRDye 680RD antibody (1:15000, LI-COR #925-68072). Fluorescent signal was captured using the Odyssey Imaging System (ThermoFisher). Quantifications of protein bands were performed using Image Studio Lite. RUNX2, MMP13, and CXCL14 levels were normalized to β-actin.

### Doxycycline treatment

Stock solutions of doxycycline hyclate (Acros Organics, 446060250) were made to a final concentration of 1 mg/ml in water, filter sterilized, and stored at −20 °C as single use aliquots. DF-1 cells and mandibles were treated in culture with the stock solution diluted in DMEM, with minocycline microspheres (Arrestin) added directly to each well, or by suspending microspheres in PBS and injecting them into the lower jaw with a 30-gauge needle. Pluronic F-127 (Sigma, P2443-250G) was dissolved at a final concentration of 35% (w/v) in DMEM growth medium rocking at 4°C for 48 h. Dox was added to Pluronic F-127 for a final concentration of 500 ng/ml and injected into the lower jaw with an 18-gauge needle. For *in ovo* treatments, 2.5 μl (for chick) and 3.75 ul (for duck) of the 1mg/ml dox stock solution was diluted with 750 μl of HBSS. This solution was then gently pipetted into the egg adjacent to the embryo and allowed to diffuse.

### Imaging

DF-1 cells were imaged using a macroconfocal (Nikon AZ100 C2+). Timelapse experiments were carried out in a custom-made stage top incubator (Okolab) set to 37 °C, 95% humidity and 5% CO_2_. All DF-1 experiments were carried out in 6-well plates (Falcon, 08-772-1B) with 2 ml of DMEM. Lower jaw timelapse experiments were carried out on 6-well transwell membranes (Nunc, 140642) with 2 ml of DMEM. Brightfield and fluorescent images of duck HH24 mandibular primordia were captured on an epifluorescent stereomicroscope (Leica MZFLIII).

### Fluorescence-activated cell sorting (FACS)

DF-1 cells were washed with 2 mL of Trypsin followed by 3mL fresh wash. Trypsin activity was inhibited by adding 5 mL of DMEM with 10% FBS. Cells were pipetted and passed through 70 μm filter. Cells were sorted on FACSAriaII Flow Cytometer (BD Bioscience, San Jose, CA). For all sorts, debris and dead cells were eliminated using FSC-A and SSC-A gating, doublets were excluded via gating discrimination using FSC-H and FSC-W, and only GFP+ cells were collected.

### Statistical analysis

Statistical analysis carried out using Student’s t-test was performed (GraphPad Prism version 8.4.3, GraphPad Software, La Jolla, CA). When multiple comparisons were made, p-values were adjusted using the Benjamini–Hochberg procedure (Benjamini and Hochberg, 1995).

## Acknowledgements

We thank Tony Qu, Austen Lucena, Paul Asfour, Kate Woronowicz, and Jessye Aggleton for laboratory assistance and/or comments on the manuscript; T. Dam at AA Lab Eggs for fertilized chick and duck eggs; and the UCSF Biological Imaging Development Core (BIDC) for microscopy support. The pmScarlet-i_C1 was a gift from Dorus Gadella (Addgene, #85044). The AAVS1 Puro Tet3G 3xFLAG Twin Strep was a gift from Yannick Doyon (Addgene, # 92099). The pKanCMV-mClover3-mRuby3 was a gift from Michael Lin (Addgene, #74252). The DHB was a gift from Dave Matus. The PGK promoter was a gift from Jonathan Brunger via Tamara Alliston.

## Data Availability

All datasets and constructs will be made publicly available at the time of publication.

